# Muscle Atrophy: Different Strains of Mice Respond Differently to Single Limb Immobilization

**DOI:** 10.1101/2020.11.03.367169

**Authors:** Camilla Reina Maroni, Michael A Friedman, Yue Zhang, Michael J McClure, Stefania Fulle, Charles R. Farber, Henry J. Donahue

**Affiliations:** Department of Neuroscience, Imaging, and Clinical Sciences, University “G. d’Annunzio” Chieti-Pescara, Chieti, Italy; Institute for Engineering and Medicine and Department of Biomedical Engineering, Virginia Commonwealth University, Richmond, VA; Center for Public Health Genomics, University of Virginia, Charlottesville, VA

**Author notes:** Institute for Engineering and Medicine, Virginia Commonwealth University, Richmond, Virginia, United States of America. Corresponding author: Henry J. Donahue, Ph.D., E – mail.

**Keywords:** Immobilization, muscle atrophy, genetic variation

## Abstract

To examine whether genetic variability plays a role in muscle adaption to its mechanical environment, we examined hind limb immobilization in five different strains of mice: CAST/EiJ, NOD/ShiLtJ, NZO/HILtJ, 129S1/SvImJ and A/J. Mice had one limb immobilized by a cast for three weeks. The response to immobilization was dependent on the strain of mice examined. A/J mice lost the most body weight following immobilization and displayed a significant decrease in physical activity. None of the other strains displayed a significant decrease in activity. Food consumption was significantly increased in NOD/ShiLtJ mice. All other strains had non-significant changes in food consumption. Muscle mass/body weight was decreased by immobilization, to varying degrees, in all strains except 1291/SvImJ. Casting decreased absolute muscle mass in both quadriceps and gastrocnemius in NOD/ShiltJ and NZO/HILtJ mice, two strains that can develop diabetes, but not in the other strains. Three weeks of immobilization caused a significant increase in quadriceps levels of atrogenes in CAST/EiJ mice but not in other strains. Immobilization caused a significant increase in quadriceps and gastrocnemius levels of *Myh4* in CAST/EiJ mice but not in any other strain. A similar trend was observed for *Myh7* in gastrocnemius muscle. Immobilization resulted in a decrease the p-p70S6K1/total p706SK1 ratio in quadriceps of NOD/ShiLtJ mice and the gastrocnemius of A/J mice, but not in other strains. Immobilization did not affect the p-4EBP1/total 4EBP1 ratio in quadriceps of any of the strains examined. However, the p-4EBP1/total 4EBP1 ratio in gastrocnemius was greater in immobilized, relative to control, limbs in CAST/EiJ mice. Muscle mass normalized to body weight in both gastrocnemius and quadriceps displayed the greatest degree of heritability. These results reveal remarkable variability in responsiveness to immobilization across five different genetic strains of mice.

## 1. Introduction

The maintenance of muscle mass is critical for optimal quality of life, and mechanical loading plays an important role in maintaining muscle mass [1]. Different types of mechanical unloading, such as that resulting from bed rest, immobilization, spaceflight and decreased physical activity can result in disuse atrophy with serious consequences including increased propensity for falls, severe functional decline and eventually death [2]. Unfortunately, the mechanism underlying disuse atrophy is only partially understood.

A better understanding of the mechanism underlying unloading-induced muscle loss, or disuse atrophy, could lead to the development of novel and more effective countermeasures and therapeutics not only for disuse atrophy but also age-related sarcopenia. In humans, heritability is approximately 66% for muscle mass and 80% for lean body mass. This suggests that genetic variability may contribute to muscle mass and therefore may also contribute to muscle adaptation to unloading. The role of genetics has also been studied in mouse models. For instance, Judex *et al.* [3] identified one Quantitative Trait Loci (QTL) on chromosome 5 for unloading-induced loss in muscle cross-sectional area in the F2 offspring of a double cross between BALB/cByJ22 and C3H/HeJ23 mice. This QTL accounted for 5% (p<0.001) of the variability in muscle loss in response to unloading in the F2 mice. We are not aware of other studies that have examined the genetics of unloading-induced muscle loss. Exploring how genetic variability contributes to unloading-induced muscle loss could provide mechanistic insight into disuse atrophy.

To examine the effect of genetic variability on the response of muscle to unloading, we exposed five different strains of mice to hindlimb immobilization. The strains examined were A/J, 129S1/SvImJ, NOD/ShiLtJ, NZO/HILtJ, and CAST/EiJ. We chose to examine these strains because they are founder strains of Diversity Outbred (DO) mice. DO mice are a recently developed high-resolution mapping population that enables gene discovery for traits such as the response of muscle to unloading [4]. DO mice can be used to investigate the genetics of a wide range of complex diseases. Examining these founder strains will not only enable us to examine the role of genetic variability in the response of muscle to unloading but will also provide insight into the usefulness of DO mice in future studies.

To assess the role of genetic variability in disuse atrophy we quantified unloading-induced changes in muscle mass as well as proteins and genes critical to muscle turnover. Muscle atrophy results from an imbalance of muscle protein breakdown (MPB) and muscle protein synthesis (MPS). MPB and MPS balance is at least partially regulated by Akt which activates the protein kinase mTOR to phosphorylate p70S6K1 and 4EBP1 resulting in increased MPS. Inhibition of Akt activates the transcription factor FOXO. FOXO then activates transcriptions of the genes encoding two atrogenes, Atrogin-1 (encoded by *FbXO32)* and MuRF-1 (encoded by *Trim63),* [5] leading to MPB. [2,6,7] Thus, increasing levels of p70S6K1 and 4EBP1 reflect increases in MPS while increasing level of Atrogin −1 and Murf-1 reflect increases in MPB.

Atrophy leads to changes in fiber type composition, and how those fibers change is dependent on the type of atrophy that occurs. Unlike age-induced atrophy which results in reduced fast-twitch fibers [8,9], disuse atrophy can lead to increased fast-twitch myosin heavy chains. For instance, Shen *et al.* [10] found different expression of *Myh4,* (encodes myosin heavy chain 4 and is expressed in muscle with fast twitch fibers), and *Myh7* (encodes myosin heavy chain ß and is expressed in muscle with slow twitch fibers) in the *deltoid* and *supraspinatus* muscle. The deltoid showed an increase in *Myh7,* in response to the microgravity of space, whereas the supraspinatus displayed an increase in *Myh4.* The same increase in fast twitch fiber type was displayed in the soleus, gastrocnemius and tibialis anterior muscles. Shenkman *et al.*[11] reported an increased level of fast-twitch fibers in vastus lateralis and soleus muscles in monkeys after 12.5 days of spaceflight. These data suggest that unloading may induce a shift from slow to fast-twitch fibers.

In this study muscle atrophy was induced by immobilization of the left hind limb through casting. We hypothesized that there would be differential, strain-dependent effects of unloading on muscle mass and expression of genes and proteins associated with muscle turnover in five strains of mice – CAST/EiJ, NOD/ShiLtJ, NZO/HILtJ, 129S1/SvImJ and A/J – after three weeks of unloading using single limb immobilization.

## 2. Methods

### 2.1. Animal model

All the animal procedures were performed with the approval of the Virginia Commonwealth University Institutional Animal Care and Use Committee. We used five Diversity Outbred (DO) founder mouse strains: CAST/EiJ, NOD/ShiLtJ, NZO/HILtJ, 129S1/SvImJ and A/J. Six male mice of each strain (they were not littermates) were purchased from Jackson Laboratories (Bar Harbor, ME, USA) and arrived at 4-14 weeks of age. The mice were given 2 to 12 weeks to acclimate. At 16 weeks of age mice of each strain had their left hind limb placed in a cast. The mice were sacrificed after three weeks of casting.

### 2.2. Casting protocol

The left hind limb of the mice were casted as previously described [12]. Mice were placed under anesthesia with isoflurane (2.5%), had the left hind limb shaved using clippers, and the skin swabbed with 70% isopropanol. Surgical tape was tightly wrapped around the left hind limb from the hip to the knee. A microcentrifuge tube was then glued on top of the tape. The tube had the bottom end removed to allow air to flow into it. Right limbs were used as contralateral controls [13]. Mice were single housed and given access to food and water *ad libitum.* Casted mice were able to freely move around the cages, dragging the immobilized limb.

### 2.3 Physical Activity and Food Consumption

Physical activity and food consumption were recorded over a 24-hour period, before and after casting. Observations were made one week prior to casting surgery and two weeks after casting. To asses individual mouse physical activity, video recordings captured for 24 hours were analyzed using the OpenField Matlab function developed by Patel *et al* [14]. Food consumption was measured by weighing food in the cages before and after the 24-hour observation period.

### 2.4 RNA isolation and quantitative RT – PCR

Total RNA from muscle samples was extracted using RNeasy Fibrous Tissue kit (Qiagen, Valencia, CA, USA). The kit used a mixture of oligo (dT) and random primers to provide an unbiased representation of the 5’ and 3’ regions of the target genes for freedom in qPCR primer design. RNA was eluted from the column with RNase-free water, and 1μl was used for quantification using a NanoDrop spectrophotometer (Thermo Fisher Scientific, Waltham, MA). Total RNA was reverse transcribed using iScript gDNA Clear cDNA Synthesis kit (Bio-Rad, Hercules, CA) following the manufacturer’s instructions. RT-qPCR was performed using 500 ng of cDNA for genes encoding the atrogenes Atrogin-1 (encoded by *FbXO32)* and MuRF 1 (encoded by *Trim63), Myh4,* (fast-twitch fiber), and *Myh7* (slow-twitch fiber). All primer sequences are shown in Supplemental Table 1.

### 2.5 Western Blot Analysis

Approximately 20 mg muscle was used for protein isolation and before homogenization all fat was removed. Muscle samples were homogenized in ice-cold buffer consisting of (mmol/L):20 HEPES (pH 7.4), 2 EGTA, 50 sodium fluoride, 100 potassium chloride, 0.2 EDTA, 50 β-glycerophosphate, 1 DTT, 0.1 phenylmethane-sulphonylfluride, 1 benzamidine, and 0.5 sodium vanadate. Protein concentration was quantified using a BCA kit (Thermo Fisher scientific), and an equal amount of total protein (30 μg) per sample was subjected to standard SDS-PAGE using Mini-PROTEAN® TGX™ 4-20% 12-wells gels (Bio-Rad, Hercules, CA). Gels were electroblotted onto PVDF membranes (Bio-Rad, Hercules, CA). Western Blot analysis was performed for total and phosphorylated 4EBP1 (T37/46, Cell Signaling Technology; Boston, MA) and total and phosphorylated p70S6K1 (T389, Cell Signaling Technology; Boston, MA) with both antibodies diluted 1:1000. The secondary antibodies were diluted 1:3000. The bound immune complexes were detected using Clarity Max™ Western ECL Substrate (Bio-Rad, Hercules, CA).

### 2.6 Statistical analyses

Repeated measures two-way ANOVAs and post-hoc tests (Tukey’s or Sidak’s multiple comparisons tests) were used to test for significant differences among the strains after three weeks of casting. GraphPad Prism version 7.0 (GraphPad Software, La Jolla, CA) was used for statistical analysis. Heritability of the muscle phenotypic characteristics was also calculated as previously described [15].

## 3. Results

### Body weight, mouse activity levels and food consumption

Body weights were recorded before and after three weeks of casting (**Table 1**). All strains displayed significant weight loss. A/J mice had the highest percentage of body weight loss compared with the other strains, and this loss was significantly more than in CAST/EiJ, and NOD/ShiLtJ mice. To analyze mouse activity, we video recorded movement around the cage for a period of 24 hours before and after casting. Immobilization significantly decreased activity only in A/J mice. All other strains had non-significant, slight decreases or large increases in distance travelled. Food consumption was significantly increased in NOD/ShiLtJ mice. All other strains had non-significant increases in food consumption except NZO/HILtJ which had a non-significant decrease in food consumption.

**Table 1.** Mouse body weight, activity and food consumption. * Significantly different than CAST/EiJ and NOD/ShiLtJ, p<0.05. Values are mean±SD. Percent change is the percent change in mean

| Mouse Strain | CAST/EiJ | NOD/ShiLtJ | NZO/HILtJ | 129S1/SvImJ | A/J |
| --- | --- | --- | --- | --- | --- |
| Day 1 Weight (g) | 17.0 ± 0.9 | 30.5 ± 2.9 | 48.9 ± 4.0 | 28.4 ± 1.8 | 27.5 ± 2.0 |
| Day 21 Weight (g) | 15.8 ± 0.8 | 27.7 ± 1.1 | 42.3 ± 3.4 | 25.3 ± 1.5 | 22.9 ± 1.8 |
| % Change | -7.3% | -8.7% | -13% | -11% | -17% * |
| p-value | 0.012 | 0.015 | 0.0002 | 0.001 | 0.0002 |
| Baseline activity (km/day) | 3.01 ± 1.09 | 1.65 ± 0.88 | 1.49 ± 0.59 | 1.91 ± 0.67 | 2.34 ± 0.52 |
| Immobilized activity (km/day) | 2.69 ± 0.70 | 2.11 ± 0.66 | 1.70 ± 0.79 | 2.10 ± 0.72 | 0.90 ± 0.74 |
| % Change | -6% | +94% | +63% | +39% | -57% |
| p-value | 0.827 | 0.486 | 0.416 | 0.530 | 0.024 |
| Baseline food consumption (g/day) | 3.7 ± 2.0 | 4.0 ± 0.4 | 5.0 ± 0.5 | 4.5 ± 1.1 | 3.3 ± 0.7 |
| Immobilized food consumption (g/day) | 4.4 ± 2.2 | 4.9 ± 0.6 | 5.0 ± 1.5 | 5.7 ± 1.6 | 4.4 ± 1.3 |
| % Change | +22% | +24% | -1.6% | +33% | +43% |
| p-value | 0.149 | 0.004 | 0.886 | 0.176 | 0.175 |

### Muscle mass

We assessed muscle mass both in absolute terms and as corrected for body weight. Assessing absolute muscle mass is appropriate when evaluating the effects of unloading in one strain. This is because in each mouse the contralateral limb serves as a control for the immobilized limb in the same mouse. However, when evaluating muscle mass across strains it is necessary to normalize to body weight. There was a significant immobilization and mouse strain interaction for absolute gastrocnemius mass, absolute quadriceps mass, and quadriceps mass/body weight. There were significant main effects of mouse strain and immobilization on gastrocnemius mass/body weight. Interestingly, immobilization significantly decreased quadriceps and gastrocnemius absolute muscle mass in only NOD/ShiltJ and NZO/HILtJ mice (**Fig. 1B**). CAST/EiJ, 129S1/SvImJ and A/J quadriceps and gastrocnemius absolute muscle mass were not affected by casting. Quadriceps muscle mass/body weight was significantly decreased by immobilization, to varying degrees, in all strains except 129S1/SvImJ. Immobilization resulted in the most significant decrease in quadricep muscle mass/body weight NOD/ShiltJ mice. Gastrocnemius muscle mass/body weight was significantly decreased, to varying degrees, in all strains except 129S1/SvImJ and A/J, and again the most significant loss was in NOD/ShiltJ mice. Finally, control muscle mass/body weight varied across strains (**Fig. 1B**). Quadriceps mass/body weight was significantly different between 5 strain pairs: CAST/EiJ vs. NOD/ShiLtJ (p=0.007); CAST/EiJ vs. A/J (0.036); NOD/ShiLtJ vs. NZO/HILtJ (p<0.0001); NZO/HlLtJ vs. 129S1/SvImJ (p=0.037) and NZO/HILtJ vs. A/J (p=0.0003). However, gastrocnemius mass/body weight was significantly different in only one pair, CAST/Eij vs. NZO/HILtj (p=0.031).

**Fig. 1.**
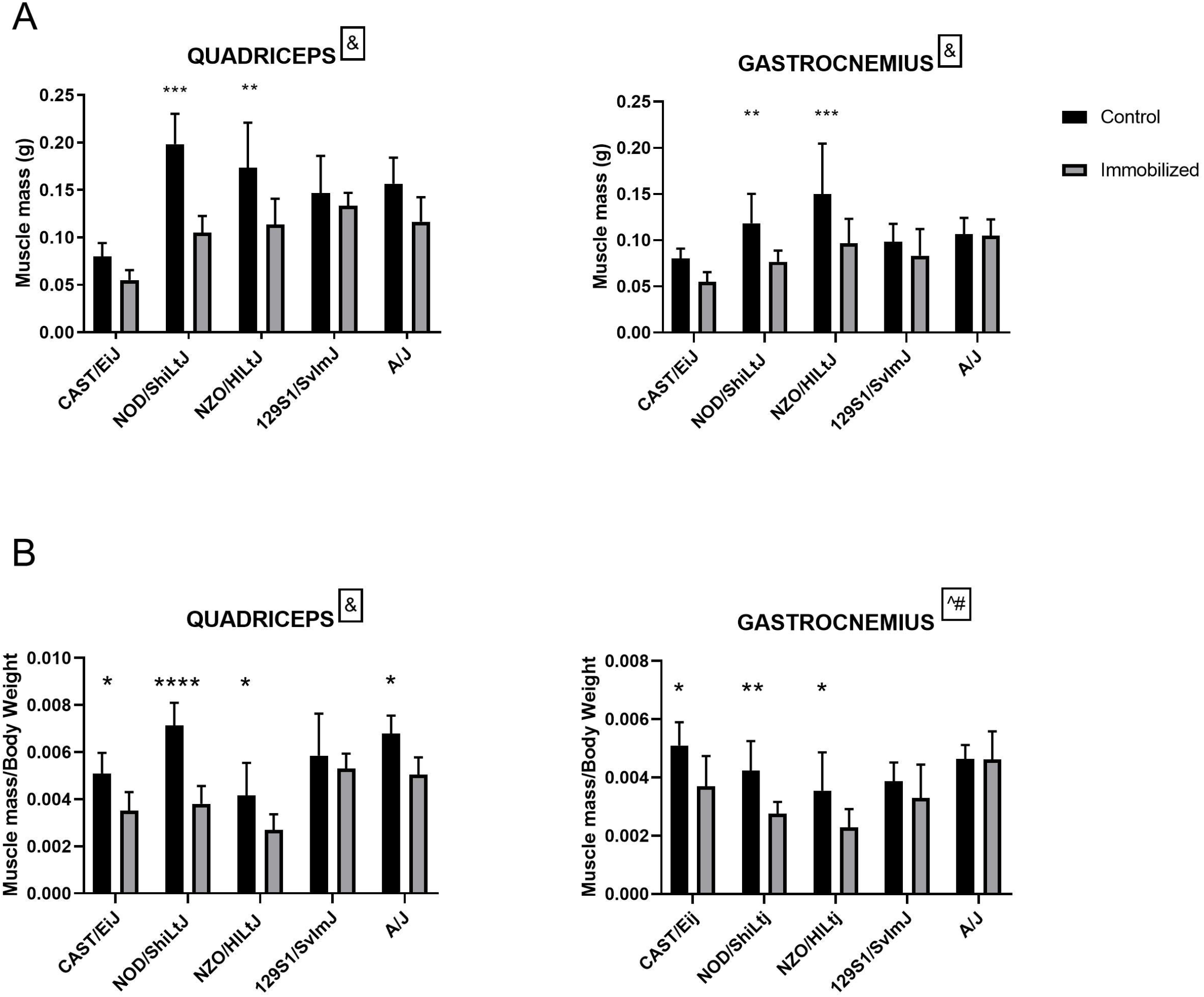
Effect of three weeks of casting on quadriceps and gastrocnemius muscle mass (A) and muscle mass normalized to body weight (B). Results for grey bars are data from control mice and black bars are data from mice exposed to immobilization by casting. Values are means±SD; n=6 for immobilized and control samples for five strains of mice. Numbers above bars are the fold changes (immobilized/control) for each strain. & Significant immobilization and mouse strain interaction (p<0.05); ^Significant main effect of mouse strain (p<0.05); #Significant main effect of immobilization; *p<0.05; **p<0.01; ***p<0.001; ****p<0.0001.

### Atrogene levels

To determine the level of muscle atrophy induced by casting, the levels of the atrogenes, *FbXO32* (encodes Atrogin-1) and *Trim63* (encodes MuRF1), were quantified by RT-qPCR (**Fig. 2**). There was a significant immobilization and mouse strain interaction for quadriceps and gastrocnemius levels of *FbXO32* and gastrocnemius levels of *Trim63.* There was a significant main effect of mouse strain on quadriceps levels of *Trim63.* Three weeks of immobilization resulted in a significant increase in quadriceps and gastrocnemius levels of *FbXO32* and *Trim63* in CAST/EiJ mice but not in any other strain. Control levels *FbXO32* and *Trim63* in both quadriceps and gastrocnemius were similar across all strains.

**Fig. 2.**
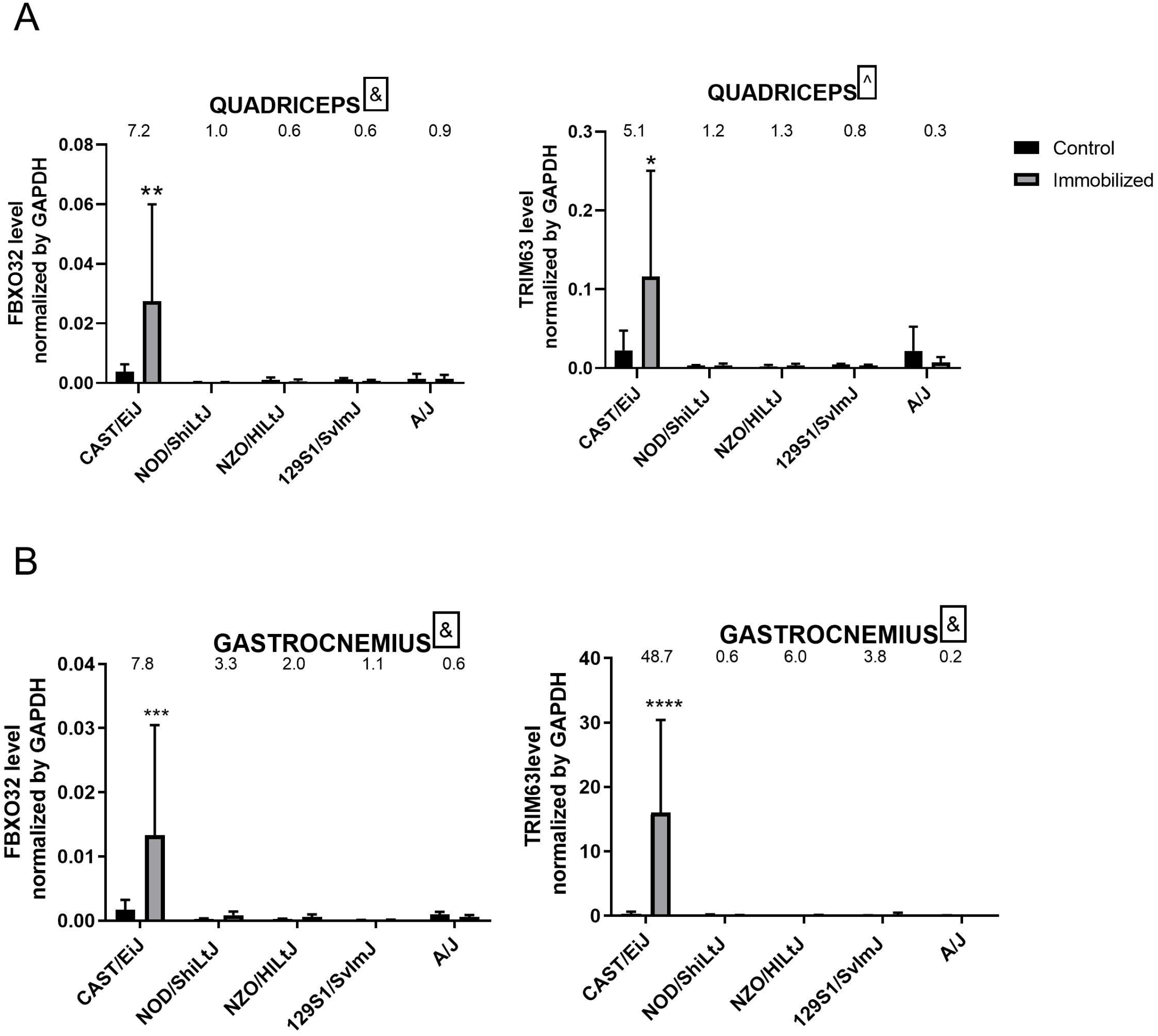
Effect of three weeks of casting on Atrogenes (FbXO32 and Trim63) expression levels in (A) quadriceps and (B) gastrocnemius muscles. Values are means±SD; n=6 for immobilized and control samples for five strains of mice. Grey bars are data from control limbs, and black bars are data from limbs exposed to immobilization by casting. Numbers above bars are the fold changes (immobilized/control) for each strain. ^Significant immobilization and mouse strain interaction (p<0.05); ^Significant main effect of mouse strain (p<0.05); *p<0.05; **p<0.01; ***p<0.001; ****p<0.0001; immobilized vs control within the same strain.

### Muscle fiber type gene levels

As previously mentioned, unloading also affects muscle fiber type composition. Levels of mRNA for genes associated with fast-twitch fibers, *Myh4,* and slow-twitch fibers, *Myh7,* were quantified (**Fig. 3**). There was a significant immobilization and mouse strain interaction for quadriceps and gastrocnemius levels of *Myh4* and gastrocnemius levels of *Myh7.* There was a significant main effect of mouse strain on quadriceps levels of *Myh7.* Three weeks of immobilization resulted a significant increase in quadriceps and gastrocnemius levels of *Myh4* in CAST/EiJ mice but not in any other strain. A similar trend was observed for *Myh7* in gastrocnemius muscle. However, *Myh 7* levels were not affected by immobilization in quadraceps of any strain. Control levels *Myh4* and *Myh7 FbXO32* in both quadriceps and gastrocnemius were similar across all strains.

**Figure 3.**
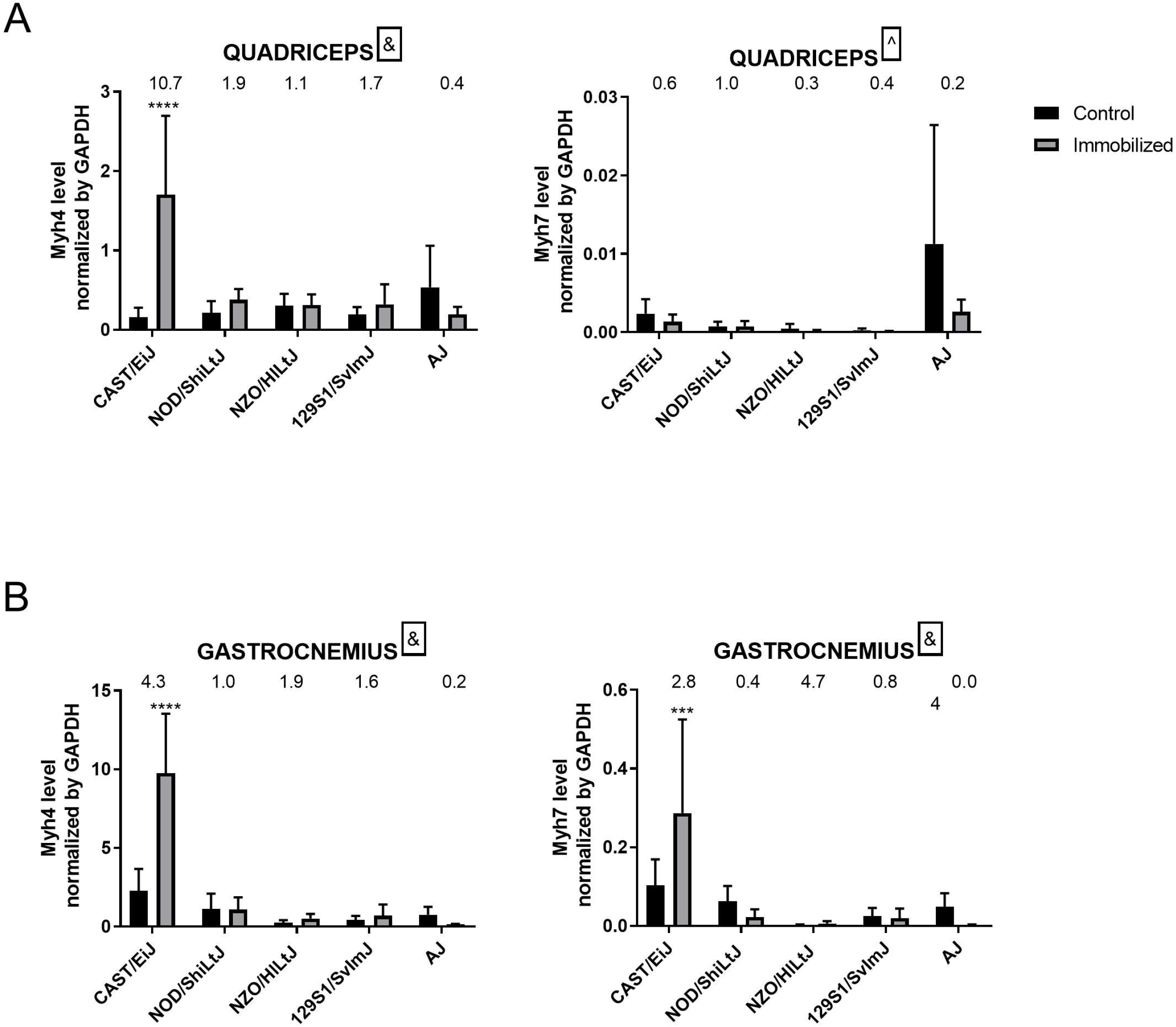
Effect of three weeks of casting on fiber type composition (Myh4 – fast twitch fiber and Myh7-slow twitch fiber) gene levels in (A) quadriceps and (B) gastrocnemius muscles in five strain of mice. Grey bars are data from control mice and black bars are data from mice exposed to immobilization by casting. Values are means±SD; n=6 for immobilized and control samples for five strains of mice. Numbers above bars are the fold changes (immobilized/control) for each strain. ^Significant immobilization and mouse strain interaction (p<0.05); ^Significant main effect of mouse strain (p<0.05); ****p<0.0001; immobilize vs control within same strain.

### Protein synthesis markers

The mTOR pathway is related to the maintenance of MPS. A central regulator of protein synthesis in skeletal muscle is mTORC1. Previous human and animal studies demostrated that the inibition of mTORC1 by rapamycin associated with excercise did not significantly reduce the protein synthesis rate [16–18]. However, by quantifying phosphorylation of p70S6K1 and 4EBP1, we indirectly assessed mTOR activity in quadriceps and gastrocnemius muscles.

There was a significant immobilization and mouse strain interaction for quadriceps p-p70S6K1/total p706SK1 ratio. There was a significant main effect of immobilization on quadriceps p-4EBP1/total 4EBP1 ratio. No other main effects were significant for p-p70S6K1/total p706SK1 and p-4EBP1/total 4EBP1. Three weeks of immobilization resulted in a significant decrease the p-p70S6K1/total p706SK1 ratio in quadriceps of NOD/ShiLtJ mice (**Fig. 4A**) and the gastrocnemius of A/J mice (**Fig. 4A**). Immobilizing did not affect the p-p70S6K1/total p706SK1 in quadriceps or gastrocnemius of any other strain. Immobilization did not affect the p-4EBP1/total 4EBP1 ratio in quadriceps of any of the strains examined (**Fig. 4A**). However, the p-4EBP1/total 4EBP1 ratio in gastrocnemius was significantly greater in immobilized, relative to control, limbs in CAST/EiJ mice (**Fig. 4B**). The p-p70S6K1/total p706SK1 ratio in control quadriceps was similar in all strains, except NOD/ShiLtJ in which levels were significantly greater than all other strains (NOD/ShiLtJ vs. CAST/EiJ, p=0.0015; NOD/ShiLtJ vs. NZO/HILtJ, p=0.0028; NOD/ShiLtJ vs. 129S1/SvImJ, p=0.0018; NOD/ShiLtJ vs. A/J, p=0.0009). The p-p70S6K1/total p706SK1 ratio in control gastrocnemius was significantly greater in A/J mice relative to other strains (CAST/EiJ vs. A/J, p=0.0174; NOD/ShiLtJ vs. A/J, p=0.0175; NZO/HILtJ vs. A/J, p=0.0442; 129S1/SvImJ vs. A/J, p=0.0133). The p-4EBP1/total 4EBP1 ratio in in control quadriceps was similar in all strains examined. However, the p-4EBP1/total 4EBP1 ratio in control gastrocnemius was significantly reduced in A/J mice relative to the other strains examined.

**Fig. 4.**
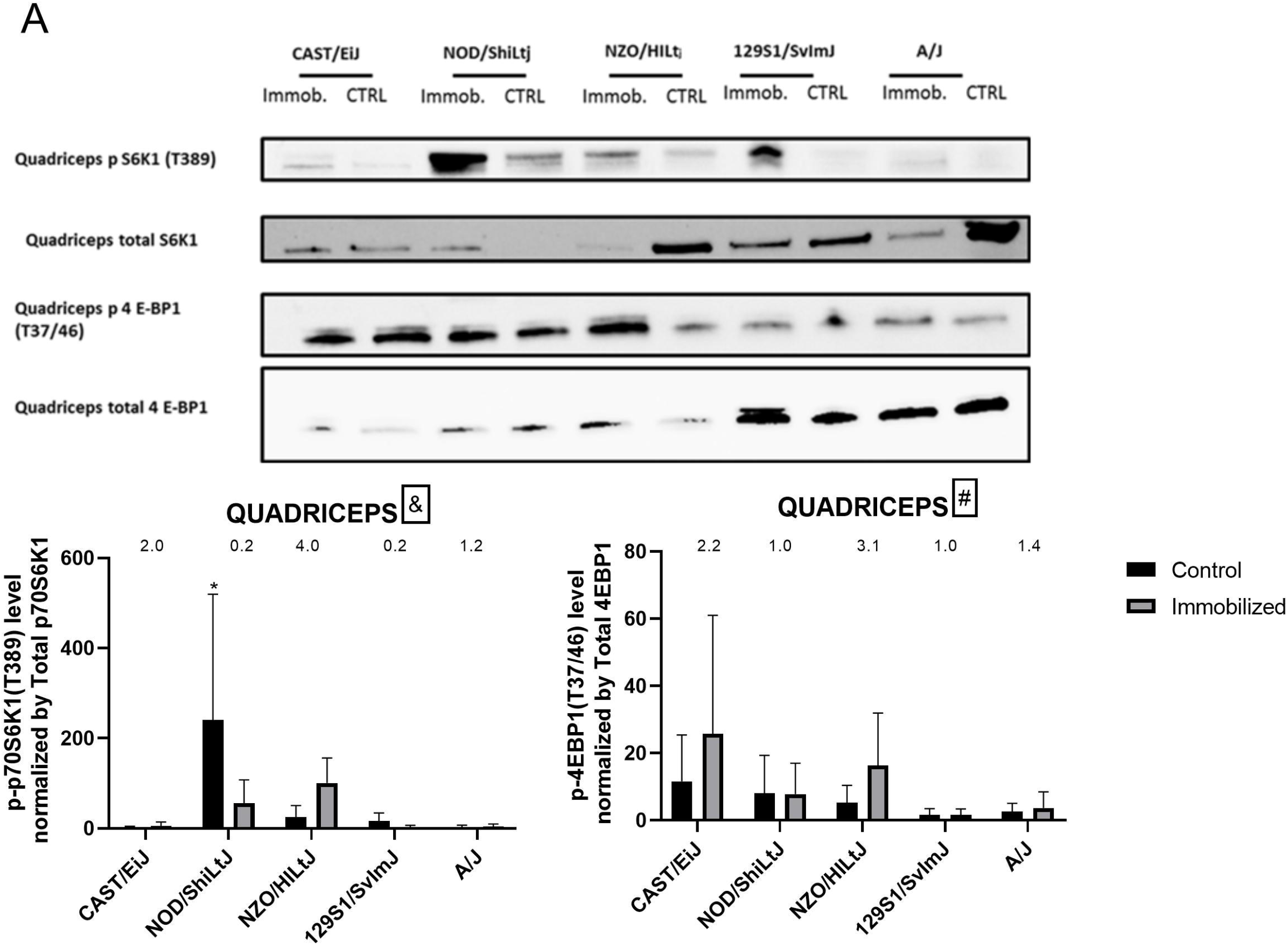
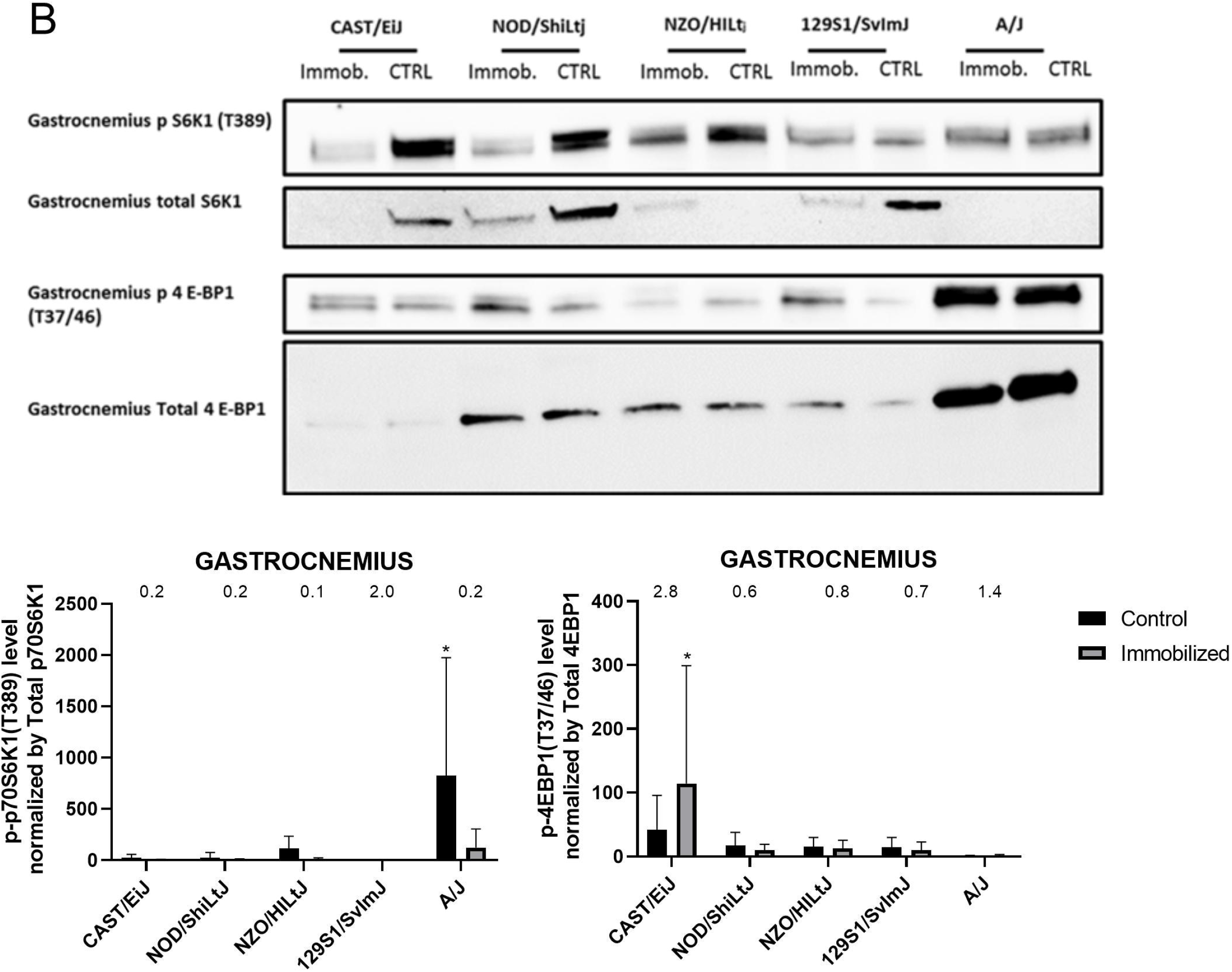
Effect of three weeks of casting on phosphorylation of p70S6K1 and 4E – BP1 in (A) quadriceps and (B) gastrocnemius muscles. All western blots were quantified and bar graphs represent mean ± SD; n=6 for immobilized and control samples for five strains of mice. Grey bars are data from control limbs, and black bars are data from limbs exposed to immobilization by casting. Numbers above bars are the fold changes (immobilized/control) for each strain ^Significant immobilization and mouse strain interaction (p<0.05); #Significant main effect of immobilization (p<0.05); * p<0.05 immobilized vs control within the same strain.

### Heritability

Considering that genetics may play an important role in the response to unloading, heritability was calculated for each property analyzed. The analysis was done for each muscle (**Table 2**). Genetics had a greater impact on muscle mass/body weight than other traits. Heritability of muscle mass, *FBXO32, Myh4 and Myh7* was greatest in the gastrocnemius. On the contrary, heritability calculated for muscle mass/body weight, *Trim63,* p-p70S6K1/total p706SK1 and 4EBP1/total 4EBP1 was higher in the quadriceps than the gastrocnemius. Phosphorylated p-p70S6K1/total p706SK1 and 4EBP1/total 4EBP1 were the traits least affected by genetics.

**Table 2.** Heritability of different properties.

| Property | Quadriceps | Gastrocnemius |
| --- | --- | --- |
| Body weight | 0.4497 | 0.4497 |
| Muscle mass | 0.2299 | 0.3133 |
| Muscle mass/BW | 0.6945 | 0.5074 |
| FBXO32 | 0.2560 | 0.4749 |
| Trim63 | 0.3898 | 0.2704 |
| Myh4 | 0.2400 | 0.3697 |
| Myh7 | 0.2807 | 0.3562 |
| p-p70S6K1 | 0.1708 | 0.1490 |
| p-4EBP1 | 0.07387 | 0.02344 |

## 4. Discussion

Our results reveal remarkable variability in responsiveness to immobilization across the five different genetic strains of mice. NOD/ShiLtJ and NZO/HILtJ were the only strains that displayed muscle loss in response to unloading. These two strains displayed similar changes in activity level, lost similar amounts of body weight following immobilization and neither of these two strains displayed an increase in atrogenes or changes in muscle fiber type following immobilization. NOD/ShiLtJ mice displayed a decrease in the p-p70S6K1/total p706SK1 ratio in the quadriceps of the immobilized limbs. Interestingly, the p-4EBP1/total 4EBP1 ratio levels were not affected by immobilization in either NOD/ShiLtJ or NZO/HILtJ mice. These results are consistent with previous data from our group showing p70S6K1 changes precede 4EBP1 changes following unloading induced by hind limb suspension of C57BL/6J mice [19]. The decrease in p-p70S6K1/total p706SK1 ratio, which is the downstream target of mTOR kinase, in NOD/ShiLtJ mice suggests that changes mTOR kinase activity and decreased protein synthesis contribute to disuse atrophy in this strain.

Our finding that only NOD/ShiLtJ and NZO/HILtJ mice displayed muscle atrophy in the immobilized limb is especially interesting in light of the fact that males of both of these strains spontaneously develop diabetes [20] while this has not been reported in the other strains examined in this study. Disuse muscle atrophy is exacerbated in diabetes [21] suggesting that susceptibility of NOD/ShiLtJ and NZO/HILtJ mice to disuse atrophy may be related to their diabetic phenotype.

CAST/EiJ mice displayed the least body weight loss during the experiment, were the most physically active and displayed an intermediate muscle mass to body weight ratio in control limbs. Interestingly, CAST/EiJ mice were the only mice that displayed changes in levels of atrogenes, *Myh4* or *Myh7. Myh4* levels were increased in both the gastrocnemius and quadriceps of immobilized limbs while *Myh7* levels were increased in the gastrocnemius of immobilized limbs. These results suggest an unloading-induced switch from slow-to fast-twitch fiber type, at least in the quadriceps. This is consistent with previous studies which revealed a switch from slow to fast fiber type associated with loss of muscle mass and strength [22–28]. Additionally, unloading from exposure to microgravity results in a switch from slow to fast fiber type [8].

Despite being resistant to disuse-induced muscle atrophy, CAST/EiJ mice displayed an increase in both atrogenes in both quadriceps and gastrocnemius muscles in the immobilized limb. This was accompanied by increases in p-4EBP1, a marker of protein synthesis, in gastrocnemius expose to immobilization. These results suggest that, in response to immobilization, CAST/EiJ muscle display increased muscle protein breakdown that is compensated for by an increase in protein synthesis, resulting in no net muscle loss.

A/J mice lost the most body weight throughout the experiment and were the only strain that displayed decreased physical activity during casting. The reason for this is unclear. However, since the A/J mice had the lowest physical activity after casting, it is possible that effectively both limbs were immobilized. The decrease in activity may have caused control limb atrophy during the first 7-14 days. Then the muscles reached homeostasis between days 15-21. Despite displaying a decrease in p-p70S6K1/total p706SK1 ratio in the gastrocnemius of the immobilized limb, A/J mice appeared, at least in the quadriceps, resistant to immobilization-induced muscle atrophy. This suggests that increases in atrogene expression, which did not change in A/J mice, are necessary but perhaps not sufficient, for immobilization-induced muscle atrophy. 129S1/SvImJ mice were resistant to immobilization-induced muscle atrophy. They lost a modest amount of body weight during the experiment but displayed no changes in physical activity as a result of casting. The immobilized limbs had similar levels, relative to control limbs, of markers of fast- and slow-twitch muscle fibers, atrogene levels, and markers of protein synthesis.

To better understand the role of genetics in the response to unloading, heritability for each property was calculated based on the percent difference between immobilized and control limbs. The results revealed a variety of impacts of genetics on these properties. Muscle mass normalized to body weight in both gastrocnemius and quadriceps displayed the greatest degree of heritability suggesting that this phenotype is the most affected by genetic variability. Interestingly the two markers of protein synthesis p-4EBP1/total 4EBP1 ratio and p-p70S6K1/total p706SK1 ratio were influenced the least by genetic variability. This suggests a disconnect between muscle mass/body weight and protein synthesis, at least as regards the effects of genetic variability. We are unaware of literature reports of the effect of disuse with casting in these strains of mice.

This study reveals differences in the response to unloading among the five mouse strains and suggests that DO mice, derived from these founders and three additional strains, would be powerful tools in identifying the mechanisms underlying disuse atrophy. However, this study has limitations that can be addressed with future studies. For instance, an examination of which genes are responsible for protecting muscle from unloading in the strains that showed minimal response to unloading would be insightful. This could be accomplished using RNA sequencing of muscle tissue samples. Additionally, our finding that immobilization did not reduce muscle mass in every strain but did affect at least one endpoint in all strains except 129S1/SvImJ, suggests that the duration of exposure to immobilization may differentially affect strain-dependent response to immobilization. Another limitation is that we used only male mice and it is well known that sex differences exist between male and female mice. Therefore, one of the next steps could be to study the response to unloading in female mice of the DO founder strains.

## 5. CONCLUSIONS

This study is the first to report a differential muscle response of five DO founder strains of mice to immobilization. The results confirm our hypothesis that genetic variability plays an important role in the development of muscle atrophy, leading to differential, strain-dependent effects of unloading on muscles in these strains of mice. Additionally, we now have the basis for using DO mice in order to identify specific gene changes that protect muscle from disuse atrophy. In this way, a therapeutic strategy to limit the development of unloading-induced sarcopenia could be developed.

## ACKNOWLEDGMENTS

This work was funded by NASA grant 80NSSC18K1473.

## REFERENCES

1. de Boer MD, Seynnes OR, di Prampero PE, Pišot R, Mekjaviċ IB, Biolo G, et al. Effect of 5 weeks horizontal bed rest on human muscle thickness and architecture of weight bearing and non-weight bearing muscles. European Journal of Applied Physiology. 2008. pp. 401–407. doi:10.1007/s00421-008-0703-0

2. Atherton PJ, Greenhaff PL, Phillips SM, Bodine SC, Adams CM, Lang CH. Control of skeletal muscle atrophy in response to disuse: Clinical/preclinical contentions and fallacies of evidence. American Journal of Physiology - Endocrinology and Metabolism. American Physiological Society; 2016. pp. E594–E604. doi:10.1152/ajpendo.00257.2016

3. Judex S, Zhang W, Donahue LR, Ozcivici E. Genetic and tissue level muscle-bone interactions during unloading and reambulation. J Musculoskelet Neuronal Interact. 2016;16: 174–82. Available: http://www.ncbi.nlm.nih.gov/pubmed/27609032

4. Svenson KL, Gatti DM, Valdar W, Welsh CE, Cheng R, Chesler EJ, et al. High-resolution genetic mapping using the mouse Diversity Outbred population. Genetics. 2012;190: 437–447. doi:10.1534/genetics.111.132597

5. Bodine SC, Baehr LM. Skeletal muscle atrophy and the E3 ubiquitin ligases MuRF1 and MAFbx/atrogin-1. Am J Physiol Metab. 2014. doi:10.1152/ajpendo.00204.2014

6. Schiaffino S, Dyar KA, Ciciliot S, Blaauw B, Sandri M. Mechanisms regulating skeletal muscle growth and atrophy. FEBS J. 2013;280: 4294–4314. doi:10.1111/febs.12253

7. Malavaki CJ, Sakkas GK, Mitrou GI, Kalyva A, Stefanidis I, Myburgh KH, et al. Skeletal muscle atrophy: disease-induced mechanisms may mask disuse atrophy. Journal of Muscle Research and Cell Motility. 2015. doi:10.1007/s10974-015-9439-8

8. Dedkov EI, Kostrominova TY, Borisov AB, Carlson BM. MyoD and myogenin protein expression in skeletal muscles of senile rats. Cell Tissue Res. 2003;311: 401–416. doi:10.1007/s00441-002-0686-9

9. Holloszy JO, Carlson BM. Factors Influencing the Repair and Adaptation of Muscles in Aged Individuals: Satellite Cells and Innervation. Journals Gerontol Ser A Biol Sci Med Sci. 1995;50A: 96–100. doi:10.1093/gerona/50a.special_issue.96

10. Shen H, Lim C, Schwartz AG, Andreev-Andrievskiy A, Deymier AC, Thomopoulos S. Effects of spaceflight on the muscles of the murine shoulder. FASEB J. 2017;31: 5466–5477. doi:10.1096/fj.201700320R

11. Shenkman BS. From Slow to Fast: Hypogravity-Induced Remodeling of Muscle Fiber Myosin Phenotype. Acta Naturae. 8: 47–59. Available: http://www.ncbi.nlm.nih.gov/pubmed/28050266

12. Friedman MA, Zhang Y, Wayne JS, Farber CR, Donahue HJ. Single limb immobilization model for bone loss from unloading. J Biomech. 2019. doi:10.1016/j.jbiomech.2018.11.049

13. Lang SM, Kazi AA, Hong-Brown L, Lang CH. Delayed recovery of skeletal muscle mass following hindlimb immobilization in mTOR heterozygous mice. PLoS One. 2012. doi:10.1371/journal.pone.0038910

14. Patel TP, Gullotti DM, Hernandez P, O’Brien WT, Capehart BP, Morrison B, et al. An open-source toolbox for automated phenotyping of mice in behavioral tasks. Front Behav Neurosci. 2014;8. doi:10.3389/fnbeh.2014.00349

15. Lariviere WR, Mogil JS. The genetics of pain and analgesia in laboratory animals. Methods Mol Biol. 2010;617: 261–278. doi:10.1007/978-1-60327-323-7_20

16. Drummond MJ, Fry CS, Glynn EL, Dreyer HC, Dhanani S, Timmerman KL, et al. Rapamycin administration in humans blocks the contraction-induced increase in skeletal muscle protein synthesis. J Physiol. 2009;587: 1535–1546. doi:10.1113/jphysiol.2008.163816

17. Philp A, Schenk S, Perez-Schindler J, Hamilton DL, Breen L, Laverone E, et al. Rapamycin does not prevent increases in myofibrillar or mitochondrial protein synthesis following endurance exercise. J Physiol. 2015;593: 4275–4284. doi:10.1113/JP271219

18. You JS, Mcnally RM, Jacobs BL, Privett RE, Gundermann DM, Lin KH, et al. The role of raptor in the mechanical load-induced regulation of mTOR signaling, protein synthesis, and skeletal muscle hypertrophy. FASEB J. 2019;33: 4021–4034. doi:10.1096/fj.201801653RR

19. Lloyd SA, Lang CH, Zhang Y, Paul EM, Laufenberg LJ, Lewis GS, et al. Interdependence of muscle atrophy and bone loss induced by mechanical unloading. J Bone Miner Res. 2014. doi:10.1002/jbmr.2113

20. Leiter EH. The NOD Mouse: A Model for Insulin-Dependent Diabetes Mellitus. Current Protocols in Immunology. John Wiley & Sons, Inc.; 2001. doi:10.1002/0471142735.im1509s24

21. Rudrappa SS, Wilkinson DJ, Greenhaff PL, Smith K, Idris I, Atherton PJ. Human skeletal muscle disuse atrophy: Effects on muscle protein synthesis, breakdown, and insulin resistance-A qualitative review. Frontiers in Physiology. Frontiers Media S.A.; 2016. doi:10.3389/fphys.2016.00361

22. Stein TP. Weight, muscle and bone loss during space flight: Another perspective. European Journal of Applied Physiology. 2013. pp. 2171–2181. doi:10.1007/s00421-012-2548-9

23. Fitts RH, Trappe SW, Costill DL, Gallagher PM, Creer AC, Colloton PA, et al. Prolonged space flight-induced alterations in the structure and function of human skeletal muscle fibres. J Physiol. 2010;588: 3567–3592. doi:10.1113/jphysiol.2010.188508

24. Stein TP, Wade CE. Metabolic Consequences of Muscle Disuse Atrophy. J Nutr. 2005;135: 1824S–1828S. doi:10.1093/jn/135.7.1824s

25. Rea G, Cristofaro F, Pani G, Pascucci B, Ghuge SA, Corsetto PA, et al. Microgravity-driven remodeling of the proteome reveals insights into molecular mechanisms and signal networks involved in response to the space flight environment. J Proteomics. 2016;137: 3–18. doi:10.1016/j.jprot.2015.11.005

26. Sandonà D, Desaphy J-F, Camerino GM, Bianchini E, Ciciliot S, Danieli-Betto D, et al. Adaptation of Mouse Skeletal Muscle to Long-Term Microgravity in the MDS Mission. Gimble JM, editor. PLoS One. 2012;7: e33232. doi:10.1371/journal.pone.0033232

27. Desplanches D. Structural and Functional Adaptations of Skeletal Muscle to Weightlessness. Int J Sports Med. 1997;18: S259–S264. doi:10.1055/s-2007-972722

28. Harrison BC, Allen DL, Girten B, Stodieck LS, Kostenuik PJ, Bateman TA, et al. Skeletal muscle adaptations to microgravity exposure in the mouse. J Appl Physiol. 2003;95: 2462–2470. doi:10.1152/japplphysiol.00603.2003

